# A major Role for Nrf2 transcription factors in cell transformation by KSHV encoded oncogenes

**DOI:** 10.1101/678342

**Authors:** Sapochnik Daiana, Raimondi Ana, Medina Victoria, Naipauer Julian, Mesri Enrique, Coso Omar

## Abstract

Kaposi‘s sarcoma (KS) is the most common tumor in AIDS patients and the highly vascularized patient‘s skin lesions are composed of the cells that derive from the endothelial tissue transformed by the KSHV virus. Heme oxygenase-1 (HO-1) is an enzyme upregulated by the Kaposi‘s sarcoma-associated herpesvirus (KSHV) and highly expressed in human Kaposi Sarcoma (KS) lesions. The oncogenic G protein-coupled receptor (KSHV-GPCR or vGPCR) is expressed by the viral genome in infected cells and is involved in KS development, HO-1 expression and vascular endothelial growth factor (VEGF) expression. We have characterized that vGPCR induces HO-1 expression and HO-1 dependent transformation through the Ga13 subunit of heterotrimeric G proteins and the small GTPase RhoA. We have found here several lines of evidence that support a role for Nrf2 transcription factors and family members in the vGPCR,-Ga13,-RhoA signaling pathway that converges on the HO-1 gene promoter. Our current information assigns a major role to Erk1/2 MAPK pathways as intermediate in signaling from vGPCR to Nrf2, influencing Nrf2 translocation to the cell nucleus, Nrf2 transactivation activity and consequently HO-1 expression. Experiments in nude mice show that the tumorigenic effect of vGPCR is dependent on Nrf2 suggesting this transcription factor or its associated proteins as putative pharmacological or therapeutic targets in KS.

## INTRODUCTION

Exposure of cells to environmental toxicants and potential carcinogens has been linked to pathogenesis of diseases including neurodegenerative disease, cardiovascular disease and cancer^1^. Eukaryotic cells have developed complex responses to detoxify potentially harmful substances and maintain cellular redox homeostasis in which different signaling cascades participate. The response involve the induction of cytoprotective and detoxifying enzymes consisting of phase I (cytochrome P450s) and phase II (detoxifying and antioxidant proteins) enzymes^2^. The expression of these genes attempts to restore the cell to a basal state preventing damage to cellular components sensitive to redox changes (i.e. proteins, lipids and DNA)^3^.

Heme oxygenase-1 is an inducible and ubiquitous 32-kDa enzyme that regulates heme metabolism and iron levels by catalyzing the degradation of the heme group. The products of this enzymatic reaction are carbon monoxide, free iron, and biliverdin. This last product is subsequently reduced to the antioxidant bilirubin^4^. HO-1 activity can be regulated at different levels but it depends primarily on the control of HO-1 expression at the transcriptional level^4–6^. A variety of stress-inducing stimuli, antioxidants, growth factors, and hormones^7–9^ can induce HO-1 expression.

HO-1 has been considered a cytoprotector molecule because of the antioxidant properties of the heme metabolism products. It has been involved in several physiological responses against oxidative and cellular stress and inflammation^5^. However, several studies have now expanded this notion an defined HO-1 as an important regulator of the physiology of the vasculature, vascular endothelial growth factor (VEGF) secretion, endothelial cell cycle control, proliferation, angiogenesis, and tumorigenesis^6,10–13^. A recent study shows that HO-1 expression is induced in endothelial cells infected by the angiogenic and oncogenic Kaposi sarcoma-associated herpesvirus (KSHV) and that the enzyme is highly expressed in biopsy tissues from oral AIDS-Kaposi sarcoma lesions^14^.

The most frequent type of tumor in AIDS patients is Kaposi sarcoma (KS). The multifocal angioproliferative lesions are formed by spindle cells derived from endothelial cells transformed by the KSHV^15^. It has been demonstrated that the product of the *orf* 74 from the KSHV genome, a G protein-coupled receptor (vGPCR), plays an important role in the development of KSHV-induced oncogenesis^15–18^. Only few cells in KS-like lesions express vGPCR^16^. However, down-regulation of the receptor in these cells results in a decreased expression of angiogenic factors and tumor regression^19,20^. vGPCR is homologous to the mammalian interleukin-8 receptor CXCR2^21^ but contains a mutation that allow its constitutive, ligand-independent activity. It has been shown that expression of vGPCR in fibroblasts induces transformation, angiogenesis in endothelial cells^22^ and angioproliferative lesions KS-like in mice^16,22,23^.

The transcription factor Nrf2 is a cap‘n‘collar bZip protein ubiquitously expressed and responsible for the basal and inducible expression of proteins involved in the oxidative stress response, drug metabolism and cytoprotection. Nrf2 is also linked to apoptosis, differentiation, proliferation and growth^24^. Chan et al. showed that Nrf2 is not essential for growth, development or erythropoiesis in mammalian cells but it could still have originally played this role in an avian system, due to observations made by Itoh et al. of high Nrf2 expression in chicken hematopoietic cells^25,26^. Furthermore, there is evidence of Nrf2 role in haematopoiesis in a mammalian system as it regulates the expression of HO-1 which is involved in the production of iron^27^. Using proteomic analysis to compare wild type (WT) and Nrf2 knockout (Nrf2^-^/^-^) mice, it was shown that basally, Nrf2 regulates a number of genes involved in the synthesis and metabolism of lipids^28^. The triterpenoids CDDO-Me and CDDO-Im are Nrf2 inducers and have been shown to reduce the accumulation of lipids in the livers of mice on a high fat diet via the Keap1/Nrf2 pathway^29^.

An important factor in the functioning of many transcription factors is their spatio-temporal regulation and this is no different for Nrf2; it is just as important that Nrf2 is switched on in response to a stimulus as it is that it is switched off when the stimulus has been removed. It is for this reason that is very important to highly regulate this pathway through different mechanisms responsible for preventing the aberrant activation of Nrf2. One of the most important mechanisms is proteasomal degradation, it regulates the cells response to inflammatory, hypoxic, oxidative and xenobiotic stimuli; and the Nrf2 pathway is no exception to this^30^. In unstressed conditions, the level of Nrf2 protein is maintained at very low levels by its inhibitor Keap1. This protein sequesters Nrf2 in the cytosol and facilitates its degradation via the proteasome. Under conditions of stress or in the presence of Nrf2 inducer, Nrf2 translocates to the nucleus and forms a heterodimer with small masculoaponeurotic fibrosarcoma (Maf) proteins which facilitate the binding of Nrf2 to the Antioxidant Response Element (ARE), a cis-acting enhancer sequence (TCAG/CXXXGC) in the promoter region of Nrf2-regulated genes^31,32^. The genes that are Nrf2-regulated can be classified into phase II enzymes, antioxidants, molecular chaperones, DNA repair enzymes, and anti-inflammatory response proteins^33^. The expression of these genes result in reduction of reactive compounds level such as electrophiles and free radicals to less toxic intermediates and an increasing ability of the cell to repair any damage ensued. Importantly, Nrf2 has been shown to possess an ARE sequence within its own promoter region to initiate its own transcription further enhancing the adaptive cell defense response^31^.

The vast number of compounds natural and synthetic able to induce Nrf2 can be divided into at least 10 different groups. The pharmacological activation of Nrf2 by various compounds has been proposed for use in the prevention of a number of diseases associated with oxidative stress^24^. A number of Nrf2 inducers such as sulforaphane (broccoli)^34^, curcumin (turmeric)^35,36^ and resveratrol (grapes)^37,38^ are currently in clinical trials for a variety of cancers. Interestingly there is some dispute over the exact role of Nrf2 in cancer as it seems to play a dual role potentially acting as both a tumour suppressor and an oncogenic factor. Recent observations that include mutations in the Nrf2 or Keap1 genes result in aberrant expression or ineffective regulation of Nrf2. Raised Nrf2 levels have been detected in an array of cancer tissues including lung ^39,40^ and pancreas^41,42^ and it is proposed that this provides cells with enhanced chemo-resistance as well as supporting increased proliferation thus promoting cancer growth and development^41^. Nrf2-deficient mice are more susceptible to toxicity by compounds such as paracetamol and tobacco smoke and to many diseases. Interestingly, Nrf2 ^-^/^-^ mice do survive and are able to procreate which suggests that Nrf2 is not necessarily vital for survival in unstressed cells and is only called upon in the presence of stress or insult^43^. However, Nrf2 plays a major role in health and disease and it is not surprising that Nrf2 is considered to be a potential therapeutic target. The development of a number of Nrf2 inducers as possible pharmacological agents without a complete knowledge of this pathway and its regulation highlight the need to understand and determine whether activation of Nrf2 would be beneficial in both the shortand long-term.

The protein responsible for the regulation of Nrf2 in the cytoplasm is Kelch-like ECH-associated protein 1 (Keap1), which forms a heterodímer that sequesters Nrf2 in the cytosol, inactive. Keap1 facilitates the Cul3-mediated poly-ubiquitination of Nrf2 leading to its proteasomal degradation and the association between Keap1 and the actin cytoskeleton also prevents this complex from entering the nucleus, limiting basal activity of the transcription factor^44^. Additionally, whilst Keap1 seems to be the major mechanism by which Nrf2 levels are controlled in the cell, recent research would suggest that the pathway is highly complex and supports a multi-faceted defense system. Furthermore a number of mechanisms by which Nrf2 can be regulated independently of Keap1 at the level of protein transcription, translation and by post translational modifications were described^45^.

Previous work from our laboratory and collaborators shows that vGPCR induces HO-1 mRNA and protein levels in fibroblasts and endothelial cells and that these facts correlate with increased cell proliferation, survival and VEGF-A expression, one of the determinant events in KS development. Additionally, inhibition of HO-1 expression or activity impairs the tumorigenesis induced by vGPCR in allograft tumor animal models^46^. Despite the implication of HO-1 as a vGPCR downstream target, the detailed nature of the molecular pathways that control its expression remain unknown. Several studies show that vGPCR contributes to KS development by switching on a complex network of signaling pathways, that include direct or autocrine/paracrine activation and expression of receptors, cytokines, signaling molecules, and transcription factors^17,47^. vGPCR activates downstream effectors by coupling to different subunits of heterotrimeric G proteins^48–50^. We have shown, that expression of Ga12 and Ga13 subunits mimicked vGPCR-induced HO-1 expression and transformation and that is through both Ga subunits. Our data indicate that vGPCR activates the small GTPase RhoA, a known downstream effector of Ga12/13, and that reduced expression of RhoA impairs vGPCR-induced VEGF expression and secretion, cell survival and proliferation, and transformation both in cell culture and in a murine allograft tumor model. The Ga12/13/RhoA signal transduction pathway is an important mediator of vGPCR-induced tumor growth and we have demonstrated the participation of HO-1 in this process. These findings suggest that the enzyme can be a potential therapeutic target in the treatment not only of KS but also of tumors where RhoA is oncogenic. Despite the implication of HO-1 as a vGPCR downstream target, the nature of the molecular pathways connecting the receptor to HO-1 expression remains unknown.

In this study, using a combination of biological models that include cells in culture and mice, we have shown that the effect of vGPCR signalling on HO-1 expression and tumorigenesis is mediated by ARE sites and the transcription factor Nrf2. Moreover, vGPCR not only affects the transcriptional activation of Nrf2 but it also induces Nrf2 nuclear trasnlocation. Nrf2 transcriptional activity and nuclear translocation are mainly mediated by activation of the ERK1/2 signaling pathway.

## EXPERIMENTAL PROCEDURES

### DNA Constructs

The plasmid pHO-1-Luc was provided by J. Alam and contains a 15-kb murine HO-1 promoter upstream of a luciferase gene as described previously^1^. The plasmid pCEFL-AU5-vGPCR, have been described previously^2,3^. The reporter construct pGL3pv Nrf2 3xARE Luc (minimum promoter containing 3 ARE sites in tandem) was a gift of M. Marinnisen and the pG5 Luc (minimum promoter containing 5 sites for GAL4 binding) was purchased from Promega. The expression plasmid for Nrf2 WT, pCDNA3 His B – V5 Nrf2 (Wild Type), was a kind gift from M. McMahon y J. Hayes^4^. The expression plasmid for Nrf2 Dominant Negative (Nrf2 DN) pEF/myc Nrf2 DN was provided by J. Alam^5^. The expression plasmid for vGPCR,pCEFL-AU5-vGPCR, was a kind gift from S. Gutkind^6^. pCDNA3 Gα12QL, pCEFL HA Gα13QL, pCEFL AU5 RhoAQL, pCEFL Rho N19 Dominant Negative was a kind gift from S. Gutkind^7^. The expression vectors for the MEKs were previously described^7^.

### Cell Lines and Transfections

NIH3T3 fibroblasts were maintained in Dulbecco‘s modified Eagle‘s medium (DMEM) (Invitrogen) supplemented with 10% calf serum. Stable transfections of NIH 3T3 for vGPCR were described previously^6^. Stable transfections of NIH – vGPCR for different shRNAs were performed using the Lipofectamine Plus Reagent (Invitrogen). NIH-vGPCR cells were plated at 60% confluence in 10 cm plates and transfected with 2ug of shRNA (shScramble and shRNAs targeted against Nrf2 - GIPZ Nfe2l2 shRNA Thermo RMM4532-EG18024). Transfected cells were selected with 750 ug/ml G418 (Promega Corp., Madrid, Spain) and 0.4ug/ml Puromycin (Invitrogen).

### Luciferase Reporter Assays

Cells were transfected with different expression plasmids together with 1ug of the indicated reporter plasmid per well in 6-well plates. In all cases, the total amount of plasmid DNA was adjusted with pcDNA3 empty and 0.2ug of pCDNA3-b-galactosidase. *Firefly* luciferase activity present in cellular lysates were assayed using the dual luciferase reporter system (Promega Corp.), and light emission was quantified using a luminometer (Junior Berlthold).

### Western Blot

Nrf2 and phospho-Nrf2 were detected by Western blotting with anti-Nrf2 (Abcam ab89443) and anti-phospho-Nrf2 (Abcam ab76026). For kinase analysis we used anti-ERK2 (Santa Cruz: sc-154), Anti-Phospho –ERK1/2 (Santa Cruz sc-7383), Anti-p38 (Santa Cruz sc-535-G), Anti-Phospho -p38 (Cell Signaling 9211), Anti-JNK (Santa Cruz sc-474-G), Anti-Phospho –JNK (Cell Signaling 9255S), Anti-Akt1 (Santa Cruz sc-1618), Anti-Phospho -Akt1/2/3 (Santa Cruz sc-16646-R). Proteins were visualized by enhanced chemiluminescence detection (Amersham Biosciences) using secondary antibodies coupled to horseradish peroxidase or secondary antibodies coupled to fluorophores and detected using an Odyssey System (Li-cor).

### Indirect Immunofluorescence

NIH 3T3, NIH 3T3 vGPCR, NIH 3T3 G12QL, NIH 3T3 G13QL and NIH 3T3 RhoAQL cells were seeded on glass coverslips. Cells were serum-starved for 24 h, washed twice with 1ml PBS, and then fixed and permeabilized with 4% formaldehyde and 0.05% Triton X-100 in 1ml PBS for 10 min. After washing with PBS, cells were blocked with 1% bovine serum albumin and incubated with anti Nrf2 (Abcam), as primary antibody O.N. at 4C. Following incubation, cells were washed three times with 1ml PBS and then incubated for an additional hour with the corresponding secondary antibody (1:1000) conjugated with fluorescein isothiocyanate (Molecular Probes). Cells were washed three times with 1ml PBS and stained with Propidium Iodide (1 ug/ml) (Molecular Probes) in the last wash. Coverslips were mounted in Mowiol mounting medium (Sigma) and viewed using a confocal microscope (Fluoview FV300, Olympus, Japan)

### Immunohistochemistry

Tumor, skin, and liver tissues were removed and fixed in 4% paraformaldehyde in 1_ PBS, transferred to 70% ethanol, and embedded in paraffin. Sections were hydrated in a graded xylene/ethanol series. Slides were stained with hematoxylin, dehydrated, and mounted in Glycergel mounting medium (DakoCytomation).

### Tumor Allografts in Athymic Nude Mice and Antitumor Effect of Nrf2 Silencing

NIH 3T3, NIH 3T3 vGPCR, NIH 3T3 vGPCR shScramble, NIH 3T3 vGPCR shNrf2 stable cell lines were used to induce tumor allografts in 7-week athymic (*nu/nu*) nude female mice. Cells were harvested, washed, counted, and resuspended in PBS. 1 × 10^6^ NIH 3T3 vGPCR or NIH 3T3 vGPCR shNrf2 cells were injected subcutaneously in the right flank of six and five nude mice respectively. Mice were monitored twice weekly until each animal developed one tumor in the area of the cell injection. Tumor volume and body weight were measured every other day during the period of investigation. Tumor volumes (*V*) were determined by the formula *V*_L_*W*2_0.5, with *L* being the longest cross-section and *W* the shortest. Data are mean ± S.E.M expressed as tumor volume (cm^3^) and it was calculated as described in Materials and Methods. (Two-way repeated measures ANOVA, treatment factor F_1,81_ = 2.47 p = 0.15).

### Statistical Analysis

Group differences were analized by ANOVA followed by Dunnet Test. Experimental groups are compared to the control. A *p* value <0.05 (*) was considered statistically significant.

### Image Analysis and Quantification

Different band intensities corresponding to Western blot detection of protein samples were quantified using the ImageJ software.

## RESULTS

### Elements involved in the vGPCR activation of HO-1

To determine which are the elements within the HO-1 promoter responsible for vGPCR activation we performed Luciferase Reporter Assays. For this aim we used a reporter construct of the HO-1 promoter of 4.9Kb (HO-1 4.9 Luc) and serial deletions (HO-1 3.8 Luc, HO-1 2.2 Luc, HO-1 1.4 Luc and HO-1 0.3 Luc) (Figure 1A). As shown in Figure 1B the loss of the proximal ARE reduce the transcriptional activity of HO-1 (4.9Kb) promoter. The 2.2 Luc showed an increment in the promotor activity indicating that between −3.8Kb and −2.2Kb might be a silencing sequence or a downregulation element. Altogether this result suggest that ARE sequence might be responsible for HO-1 promoter activation by vGPCR and that there are enhancer and silencing sequences within the HO-1 promoter regulated by vGPCR.

**Figure 1:**
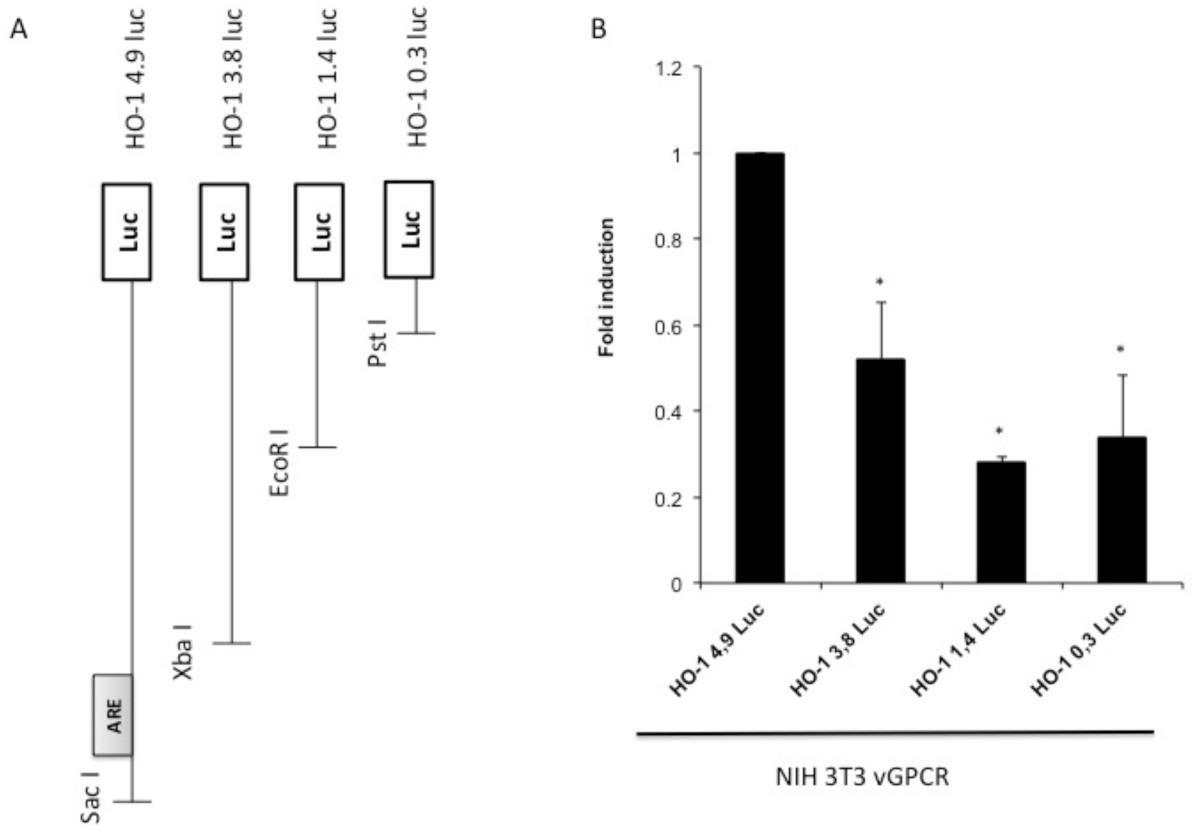
A) Serial deletions of the 4.9Kb region of the HO-1 promoter. B) Luciferase activity of serial deletions of the HO-1 promoter after co-transfection of the reporter construct with a plasmid that expresses vGPCR. The results are expressed as fold induction relative to cells transfected with the 4.9Kb promoter construct.

### Relevance of ARE sites in vGPCR activation

To determine the role of ARE sites present in the HO-1 promoter for HO-1 activation by vGPCR we peformed Luciferase Reporter Assays. We used a mínimum promoter that contains thre AREs site in tándem (3xARE Luc). We co-transfected this reporter with expression plasmids for Nrf2 WT and Nrf2 Dominant Negative (DN) (Figure 2A) as a control of responsiveness of the reporter. Nrf2 DN is a Nrf2 dominant negative that lacks the transactivation domain. Then we evaluate the effect of over-expression of vGCPR. As seen in Figure 2B, vGPCR produced an increment in the reporter activity, indicating that vGPCR may activate ARE sites in target genes. Empty pCDNA3.1 was co-trasnfected as negative control (Figure 2A column 1). To determine if this effect was mediated by the transcription factor Nrf2 we co-transfected vGPCR with plasmids that express Nrf2 WT (Figure 2C) or Nrf2 DN (Figure 2D). Interestingly co-transfection with vGPCR and Nrf2 WT produce a higher induction than the observed for vGPCR alone, suggesting that vGPCR and Nrf2 may be in the same pathway. To confirm this, we co-transfected vGPCR with Nrf2 DN. As seen in Figure 2D the induction produced by vGPCR was abolished by Nrf2 DN. Altogether, these results suggest that vGPCR activates ARE sites through Nrf2.

**Figure 2:**
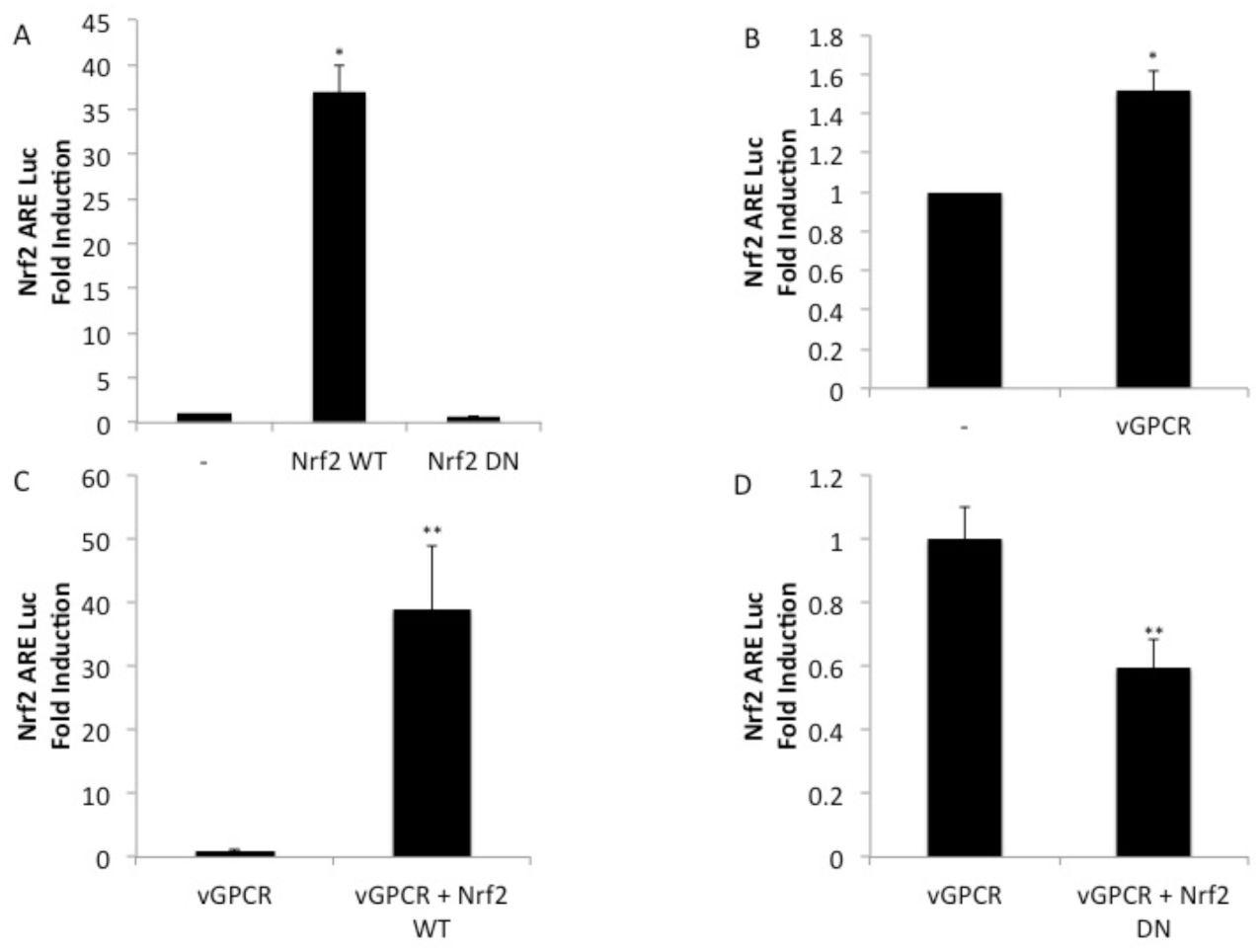
Luciferase activity of a minimum promoter with 3 ARE sites in tandem (3xARE Luc) after transfection of Nrf2 WT and Nrf2 DN as control of the system (A); effect of vGPCR (B); effect of co-transfection with vGPCR and Nrf2 WT (C) and effect of co-transfection with vGCPR and Nrf2 DN (D). The results are expressed as fold induction relative to control cells (transfected with the reporter and an empty vector).

### Rol of Nrf2 on HO-1 activation by vGPCR

To evaluate the role of Nrf2 in the induction of the HO-1 promoter by vGPCR we performed Luciferase Reporter Assay. We used a reporter with the luciferase gene upstream the murine HO-1 promoter. As seen in Figure 3A, vGCPR induce an increment in the activity of HO-1 promoter when compare to cells transfected with a control plasmid (pCDNA3.1 empty). To demonstrate that this effect was through Nrf2 we co-transfect vGPCR with Nrf2 WT (Figure 3B) or with Nrf2 DN (Figure 3C). It is shown in Figure 3B that the effect of co-transfection of vGPCR and Nrf2 WT in HO-1 promoter was bigger than the effect of vGPCR alone. More interesting, the co-transfection of vGPCR with Nrf2 DN produced a decrease of about 30% respect the effect of vGPCR alone. These results suggest that Nrf2 plays a key role in the activation of HO-1 promoter by vGPCR.

**Figure 3:**
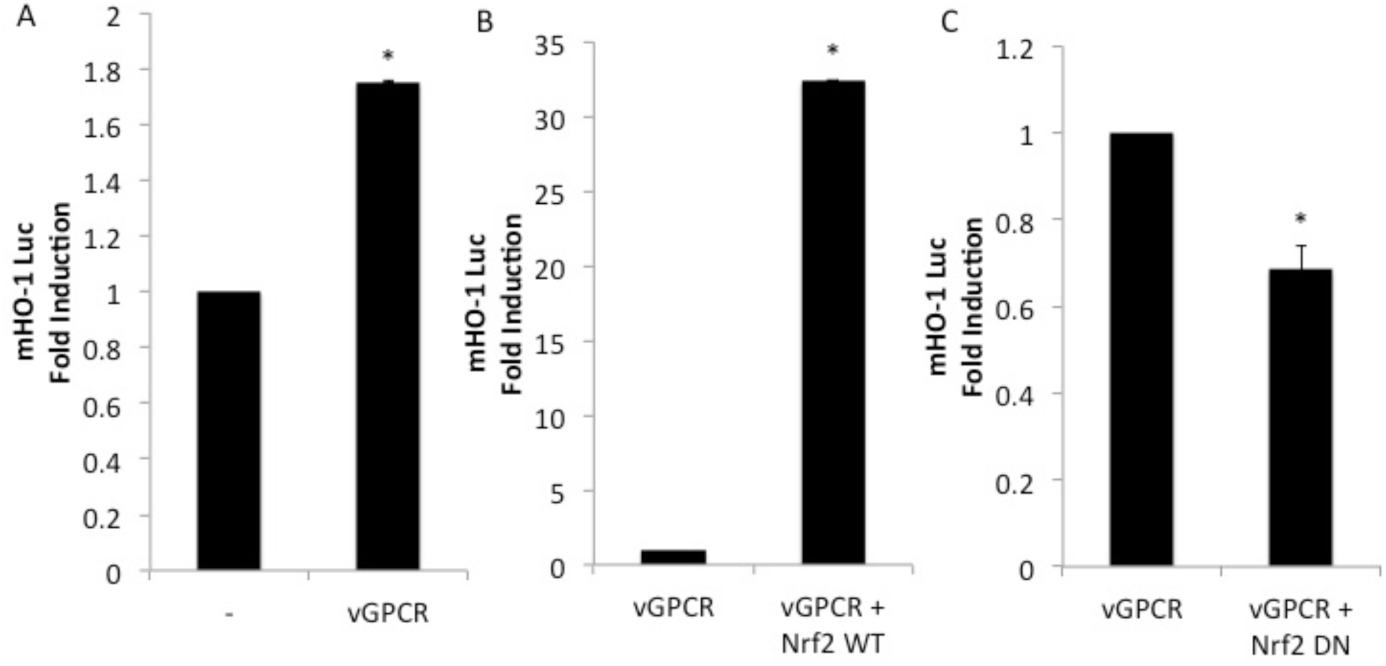
Luciferase activity of the murine HO-1 promoter after transfection with vGPCR (A); effect of co-transfection with vGPCR and Nrf2 WT (B) and effect of co-transfection with vGCPR and Nrf2 DN (C). The results are expressed as fold induction relative to control cells (transfected with the reporter and an empty vector).

### Downstream effectors involved on vGPCR activation to ARE sites

As many GPCRs, vGPCR also activate downstream effectors like Ga12, Ga13 and RhoA. To determine if those proteins where involved in the activation of the ARE sites by vGPCR we performed luciferase assay with the 3xARE Luc co-transfecting vGPCR and the downstream effectors. As shown in Figure 4A, vGPCR and the effectors downstream activate ARE sites. Moreover, the activation of ARE sites by vGPCR, Ga12 and Ga13 are RhoA dependent, as the co-transfection with RhoA Dominan Negative (RhoAN19) produce a decrease in the activation of the reporter when compared with the effect produce by vGPCR, Ga12 and G13 alone (Figure 4B). The results suggest that vGPCR, Ga12 and Ga13 activates ARE sites through RhoA.

**Figure 4:**
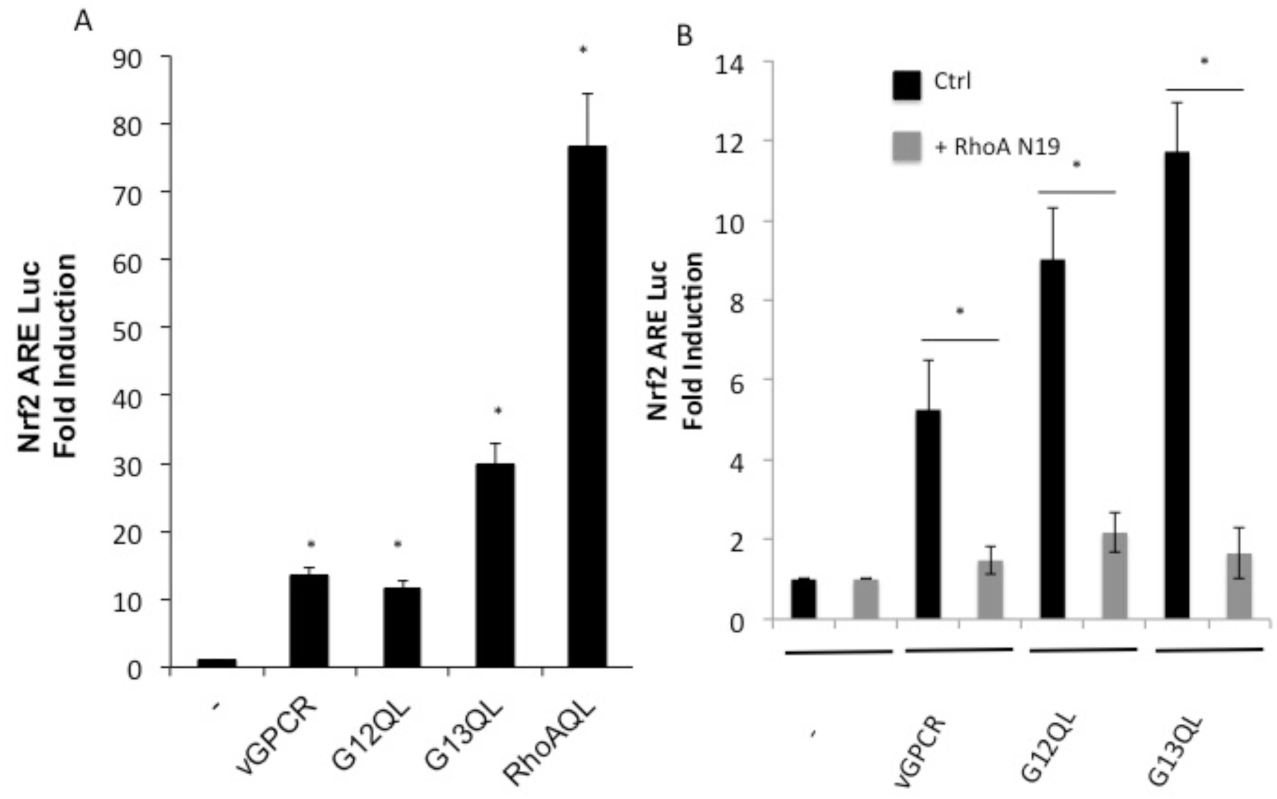
A) Luciferase activity of the 3xARE Luc reporter construct after transfection with plasmids expressing vGPCR, G12QL, G13QL and RhoAQL. The results are expressed as fold induction relative to control cells (transfected with the reporter and an empty vector). B) Luciferase activity of the 3xARE Luc after co-transfection with vGPCR, G12QL and G13QL with or without RhoAN19. The results are expressed as fold induction relative to control cells (transfected with the reporter and an empty vector).

### Effects on Nrf2 transcriptional activity

As many transcription factors, Nrf2 has to be activated on its Transcriptional Activated Domain (TAD) so it can recruit transcriptional machinery and actívate target genes transcription. To determine if vGPCR has an effect on Nrf2 TAD activation we performed luciferase assays. We co-expressed a Gal4 DBD– Nrf2 TAD fusion protein and a Gal4 Binding Element upstream luciferase and we evaluate reporter activity in control cells and in cells overexpressing vGPCR, G12QL, G13QL or RhoAQL. As shown in Figure 5 vGPCR and the elements downstream can activate this promoter suggesting that Nrf2-TAD is a target of vGCPR triggered signaling.

**Figure 5:**
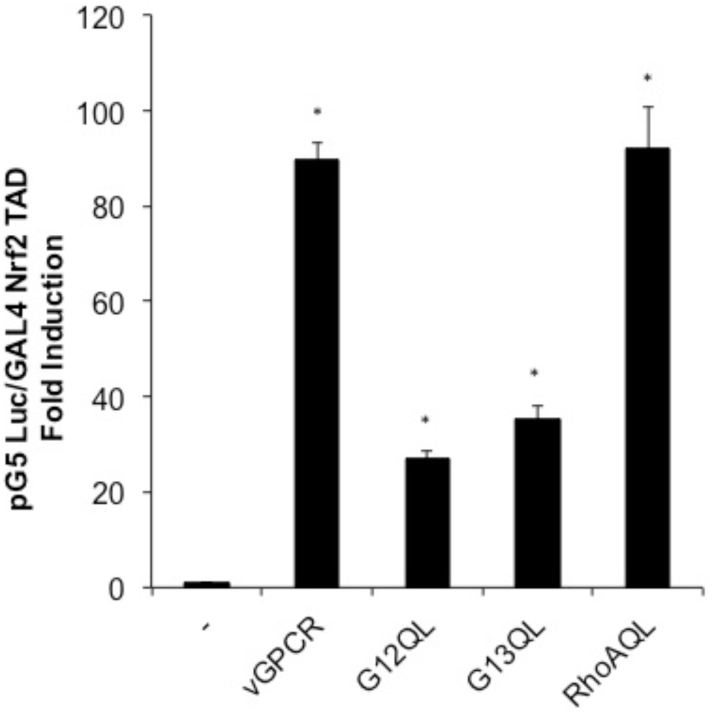
Luciferase activity of the GAL4 reporter system to study transcriptioal activation domains of transcriptios factors. pG5 Luc and pGAL4 Nrf2 TAD reporter plasmids were co-transfected with plasmids expressing vGPCR, G12QL, G13QL and RhoAQL. The results are expressed as fold induction relative to control cells (transfected with the reporter and an empty vector).

### Effects on Nrf2 nuclear translocation

As it has been shown, Nrf2 has two different subcellular localizations. In basal conditions Nrf2 localizes in the cytoplasm and its driven to proteasomal degradation by Keap1 whereas in stimulate condition the complex Nrf2-Keap1 is dissociated and Nrf2 translocates to the nucleus. To determine if vGCPR has any effect on Nrf2 nuclear translocation we performed immunofluorescence assays with specific Nrf2 antibody on NIH 3T3, NIH 3T3 vGPCR, NIH 3T3 G12QL, NIH 3T3 G13QL and NIH 3T3 RhoAQL (all stable cell lines). As shown in Figure 6 in NIH 3T3 Nrf2 has a predominantly cytoplasmic localization whereas in the 4 stable cell lines for vGPCR, G12QL, G13QL and RhoAQL, Nrf2 is found in the nucleus. Propidium Iodide staining was performed to localize nucleus. These results suggest that vGPCR and its effectors downstream have the same effect on Nrf2 nuclear translocation.

**Figure 6:**
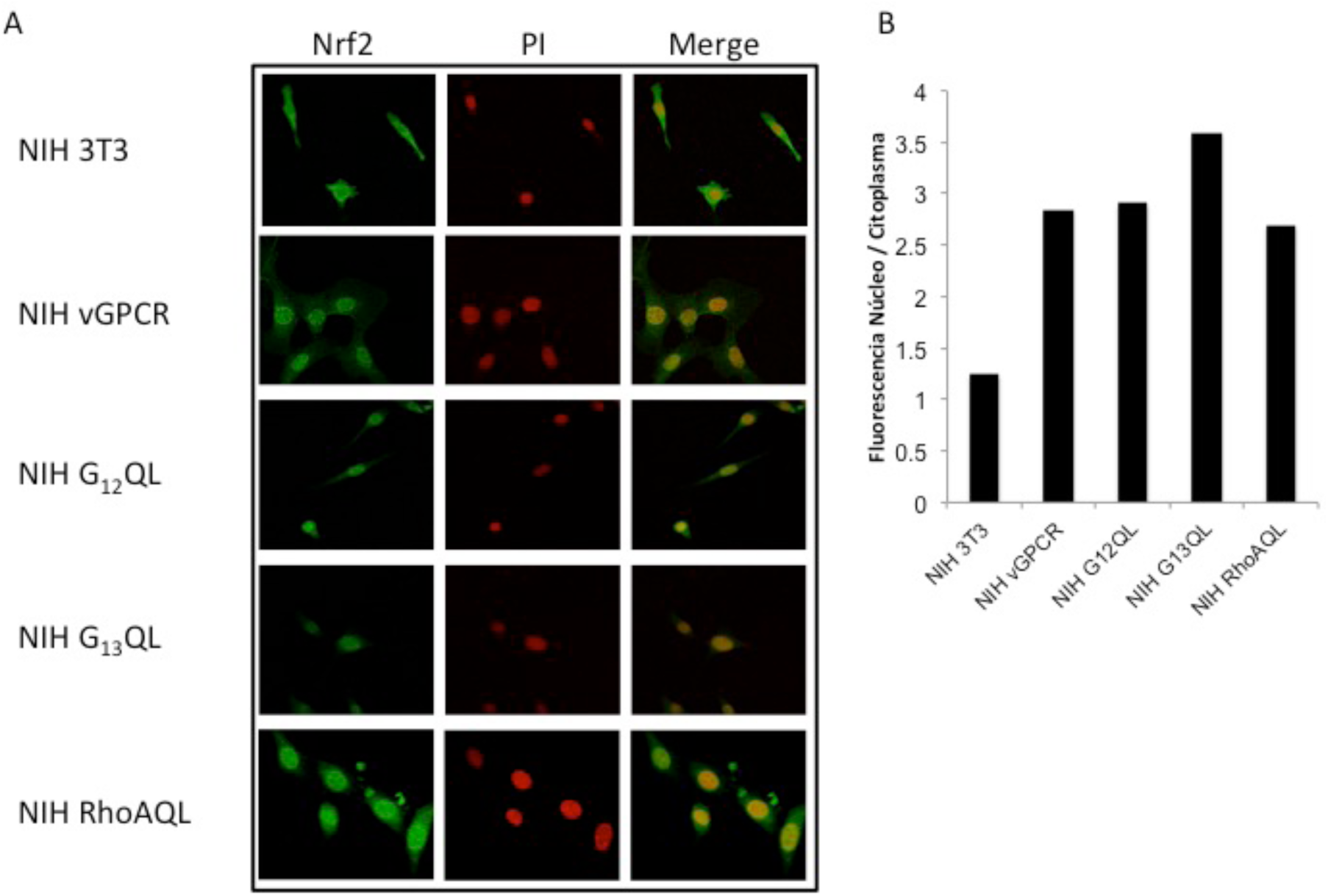
vGPCR, G12QL, G13QL and RhoAQL induced nuclear translocation of Nrf2. Confocal microscopy of NIH 3T3 stable cell lines for the expression of GPCR, G12QL, G13QL and RhoAQL were incubated with anti Nrf2 and anti rabbit FITC. For visualizing the nucleus propidium iodide was used. Magnification 40X.

### Effects of vGCPR on Nrf2 levels and Nrf2 phosphorilation

Given that it‘s been reported that Nrf2 it‘s directed to proteasomal degradation and that in stimulated conditions it can be phosphorylated we wonder if vGPCR has the ability to stabilize the protein level of Nrf2. We also wonder if vGPCR can change the phosphorylation level of Nrf2. For this purpose, we performed Western Blot using lysates of NIH 3T3 and NIH 3T3 vGPCR and evaluate the levels of Nrf2 total and Nrf2 phosphorylated in both cell lines. As seen in Figure 7, vGPCR augments the level of total Nrf2 and Nrf2 phosphorylated. These result suggest that vGPCR not only stabilize Nrf2 but it also induce Nrf2 phosphorylation.

**Figure 7:**
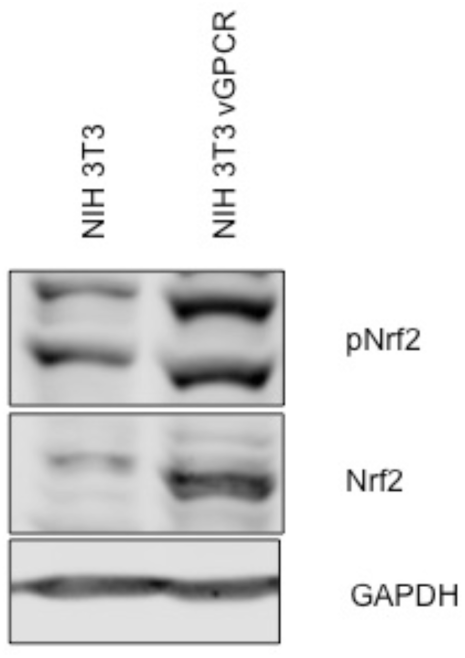
Western Blot Assays performed in NIH 3T3 and NIH 3T3 vGPCR were evaluated for Nrf2 levels and Nrf2 phosphorylation using anti-phospho Nrf2 and anti-Nrf2 as loading control we used anti-GAPDH.

### Involvement of kinases signaling pathways

It is known that Nrf2 and Keap1 can be modified by different post-tranlational modification such as phosphorylation, ubiquitination, sumoylation and many others. To determine if phosphorylation on the Nrf2-Keap1 complex is a mediator of the vGPCR effect, we evaluate different kinase activation in our model. We performed Western Blot assays with specific antibody pairs (phosphor-protein and total protein) for 4 different kinases, AKT (Figure 8A) and JNK (Figure 8B). ERK1/2 (Figure 8C), p38 (Figure 8D). In all cases, we evaluate NIH 3T3, NIH 3T3 vGPCR and a appropriate positive control (see Figure 5 A-D). As seen in Figure 8 vGPCR is capable of activate ERK1/2 and p38 but not AKT and JNK in our model.

**Figure 8:**
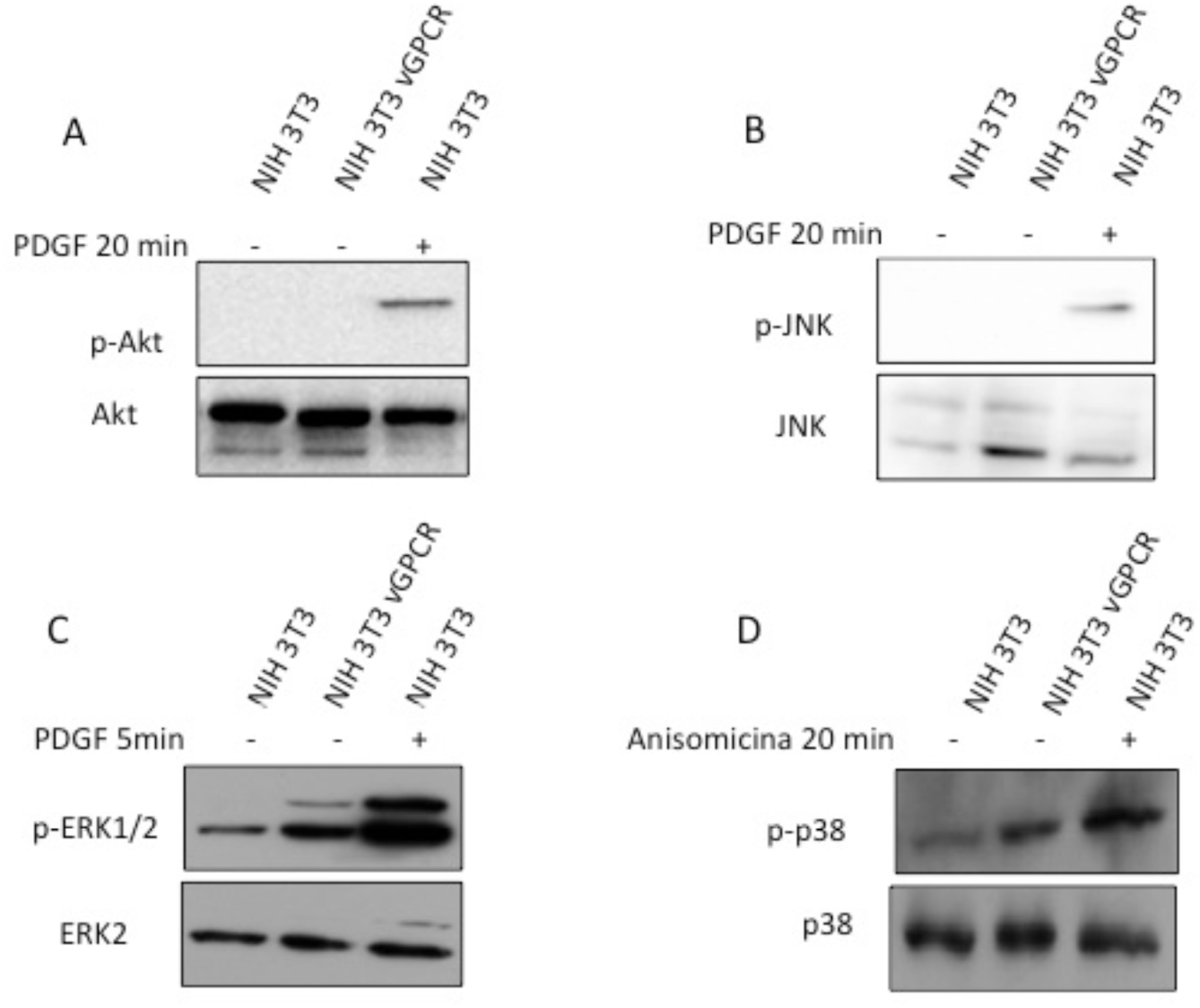
Kinase activation by vGPCR. Western Blot Assays performed in lysates of NIH 3T3 and NIH 3T3 vGPCR cells were evaluated for activation of AKT using anti-phospho AKT and anti-AKT (A) as control we used NIH 3T3 cells treated with PDGF 20 minutes; of JNK using anti-phospho JNK and anti-JNK as control we used NIH 3T3 cells treated with PDGF 20 minutes (B); of ERK using anti-phospho ERK1/2 and anti-ERK2, as control we used NIH 3T3 cells treated with PDGF 5 minutes (C); of p38 using anti-phospho p38 and anti-p38, as control we used NIH 3T3 cells treated with Anisomycin 20 minutes (D).

### vGPCR effects on Nrf2 transcriptional activity are mediated by ERK1/2

As we report previously, vGPCR has effects in Nrf2 transcriptional activity so we wanted to know if this effect was mediated by the ERK1/2 pathway. For this aim we performed Luciferase assays with the murine HO-1 promoter (Figure 9A), the 3xARE Luc (Figure 3 B) and the GAL4 – Nrf2 TAD reporter system (Figure 9C). We trasnfected NIH 3T3 vGPCR cells with the reporter construct and we activated the ERK1/2 pathway co-transfecting with expression plasmids for MEKEE (constitutive activated) or inhibited the pathway co-transfecting with MEKAA (dominant negative). In Figures 2B, 3A and 5 we showed that vGPCR induce these promoters. In Figure 9A we show that co-transfection of vGPCR and MEKEE did not produce a higher effect in HO-1 promoter activation when compare to vGPCR alone. This might be due to a saturation of the system. In the other hand, co-transfection of vGPCR with MEKAA diminished the effect observed with vGPCR alone. In Figure 9B and 9C we observed a higher effect when co-transfecting vGPCR with MEKEE than vGCPR alone. According with this result, co-transfección with vGPCR and MEKAA produced a decrease in reporter activation with respect to vGPCR. These results suggest that the ERK1/2 pathway is mediating the vGCPR effect on Nrf2 transcriptional activity and that might have a participation in HO-1 promoter activation.

**Figure 9:**
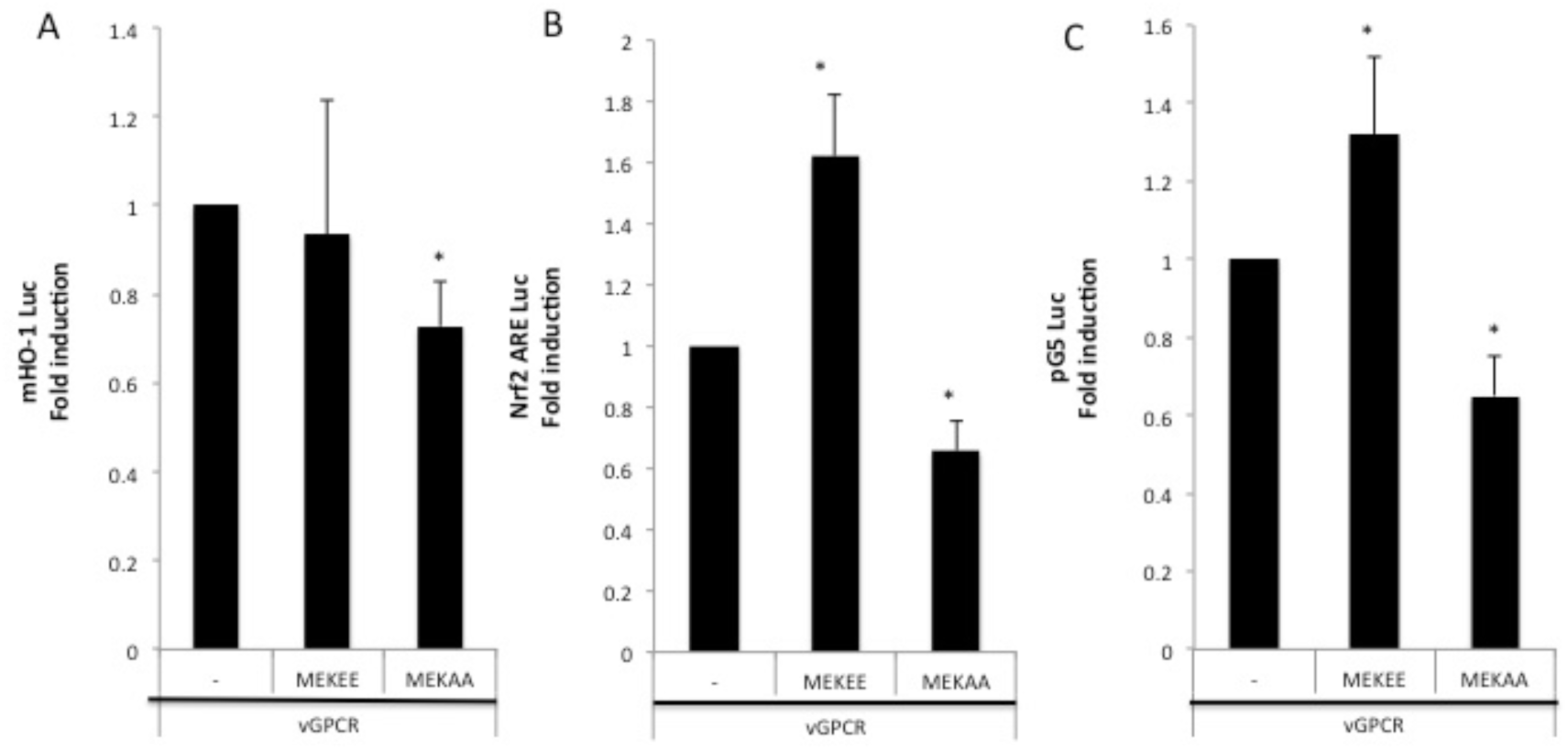
Luciferase activity of the murine HO-1 promoter (A); 3xARE Luc (B) and the GAL4 – Nrf2 TAD reporter system (C) of NIH 3T3 cells after transfection of vGPCR alone (bar 1 in each set); co-transfection of vGPCR and MEKEE (bar 2 in each set) and co-transfection of vGPCR and MEKAA (bar 3 in each set).

### vGPCR effect on Nrf2 nuclear traslocation is mediated by ERK1/2

To evaluate the ability of vGPCR to induce Nrf2 nuclear translocation by ERK1/2 signaling, we performed immunofluorescence assays using an Nrf2 specific antibody this time treating NIH 3T3 vGPCR cells with or without the MEK inhibitor PD98059 (20mM) for 2hs. Figure 10 shows that the inhibition of the ERK1/2 pathway impairs the nuclear traslocation of Nrf2. The ratio of Nuclear Fluorescence/ Cytoplasm Fluorescence diminish from 2.5 in NIH 3T3 vGCPR cells which indicates nuclear localization to a ratio of 0.5 indicative of cytoplasmic localization. This result suggest that the effect of vGPCR on Nrf2 nuclear localization is mediated by ERK1/2.

**Figure 10:**
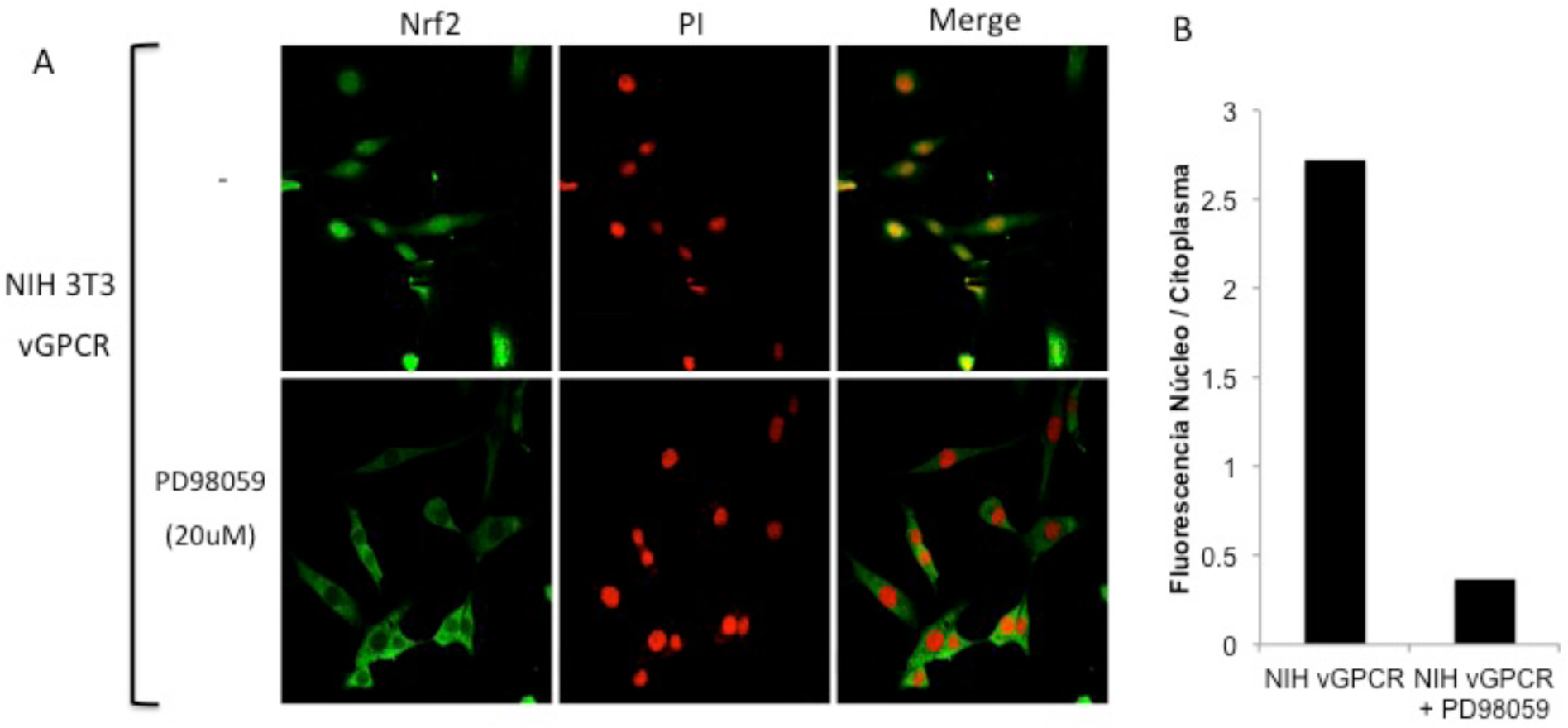
Inhibition of ERK1/2 activity block Nrf2 nuclear traslocation. Confocal microscopy of NIH 3T3 stable cell lines for vGPCR, G12QL, G13QL and RhoAQL were incubated with anti Nrf2 and anti rabbit FITC. For visualizing the nucleus propidium iodide was used. Magnification 40X.

### Rol of ERK on Nrf2 stability and phosphorilation

Whereas vGPCR increases Nrf2 stability and phosphorilation and considering that vGCPR also activates the ERK1/2 pathway we wanted to know if the ERK1/2 pathway was responsible for Nrf2 phosphorilation. For this pourpuse we treated NIH 3T3 vGPCR cells with PD98059 (20mM) for 2 hs and performed Western Blot Assays. As seen in Figure 11 the inhibition of the ERK1/2 pathway did not produce a decrease in the phosphorilation levels of Nrf2 induced by vGPCR. This result suggests that the efect of ERK1/2 may not be through direct phosphorylation of Nrf2.

**Figure 11:**
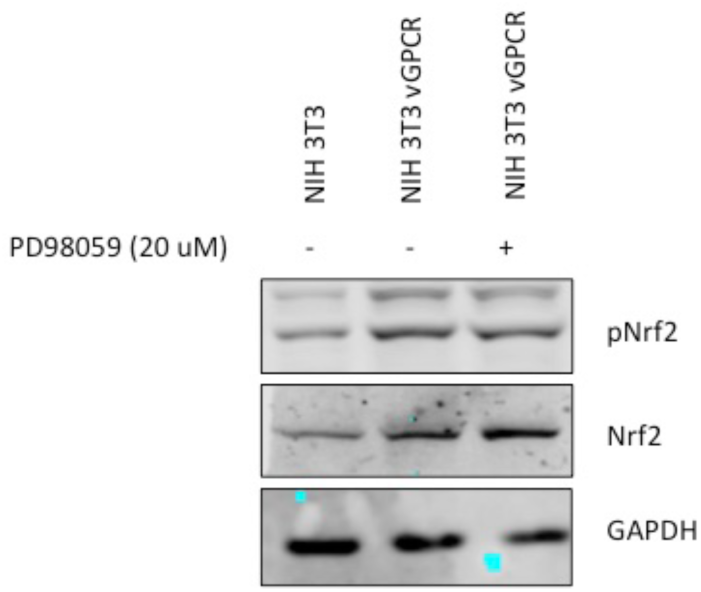
Western Blot Assays performed ising lysates of NIH 3T3, NIH 3T3 vGPCR and NIH 3T3 vGPCR cells treated with PD98059 (20mM) for 2h were evaluated for Nrf2 levels and Nrf2 phosphorylation using anti-phospho Nrf2 and anti-Nrf2 as loading control we used anti-GAPDH.

### Silencing of Nrf2 Impairs vGPCR-induced Tumorigenesis in Mice

Whereas parental NIH 3T3 cells are non-transformed, they acquire the capability to form foci in cell culture models and to induce tumors in nude mice when transformed by an oncogene. Thus, vGPCR-overexpressing NIH 3T3 cells, but not the parental cells, have been reported to induce tumors when injected into nude mice. Prompted by our findings, we used these models to investigate whether silencing Nrf2 could affect vGPCR-induced tumorigenesis *in vivo*. For this aim we transfected NIH 3T3 vGCPR cell line with different shRNAs directed against Nrf2 (NIH 3T3 vGCPR–shNrf2) and with a control shRNA (NIH 3T3 vGPCR–shScramble). We generated stable cell lines and evaluated them for Nrf2 expression levels. As shown in Figure 12A stable cell lines named NIH 3T3 vGPCR-shNrf2 2 and NIH 3T3 vGPCR-shNrf2 2.1 shown the lowest level of Nrf2 when compared with NIH 3T3 vGPCR. We injected 1×10^6^ NIH vGPCR or NIH vGPCR–shNrf2 2.1 cells into the right flank of six and five nude mice respectively and observed the mice twice a week.. In concordance with the in vitro described effect, silencing Nrf2 by shRNA has a strong impact in the tumorigenic behavior of transformed cells in vivo. Tumors produced by NIH 3T3 vGPCR – shNrf2 2.1 tend to be smaller than the ones observed in N NIH 3T3 vGPCR group (Figure 12B-C), probably due to the significant delay in tumor onset found for Nrf2 silenced cells (Figure 12D).

**Figure 12:**
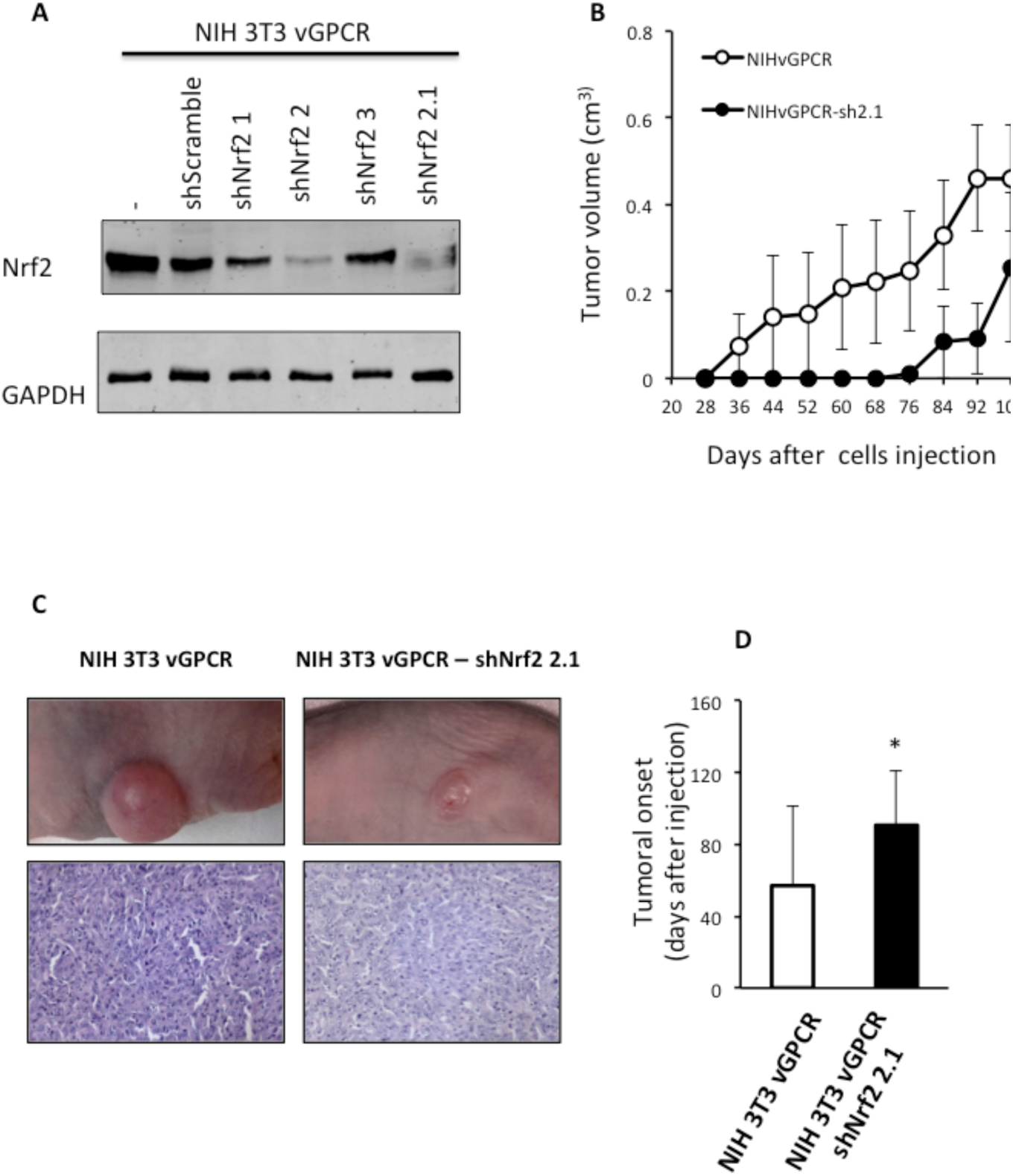
A: Western Blot Assays performed in NIH 3T3 vGPCR or different NIH 3T3 vGPCR stable cell lines for the expression of an shRNA expression plasmid targeting Nrf2. We used an anti-Nrf2 antibody for the detection of Nrf2 level in the different cell lines. B: 1 × 10^6^ NIH 3T3 vGPCR or NIH 3T3 vGPCR - shNrf2 2.1 cells were injected subcutaneously in the right flank of six and five nude mice respectively. Data are mean ± S.E.M expressed as tumor volume (cm^3^) and it was calculated as described in Materials and Methods. (Two-way repeated measures ANOVA, treatment factor F_1,81_ = 2.47 p = 0.15) C: An example of tumor-bearing mice from each group is depicted. H&E staining from representative tumors from mice injected with NIH 3T3 vGPCR or NIH 3T3 vGPCR – shNrf2 cells respectively. D: The tumor onset was defined as the first day of appearances of a measurable tumor. Data are mean ± S.E.M. NIH 3T3 vGPCR: 57± 44.33; NIH 3T3 vGPCR – shNrf2: 90.6± 30.56. Student‘s *t* test *p<0.005

## DISCUSSION

Kaposi‘s sarcoma (KS) is the most common tumor in AIDS patients and the highly vascularized patient‘s skin lesions are composed of the cells that derive from the endothelial tissue transformed by the KSHV virus^1–3^.

In previous works from our laboratory and collaborators we have contributed to show how the expression of vGPCR in fibroblasts produces transformation with foci formation in Petri plates, and tumors in nude mice. The lesions produced by these tumors in animals are rich in vessel irrigation, which resembles those of human patients with KS^1,4^. In addition, we have demonstrated that vGPCR induces the gene coding for HO-1 in both fibroblasts and endothelial cells and that this increase in HO-1 expression levels correlates and is necessary for increased proliferation and cell survival cell. On the other hand, we have shown that he inhibition of HO-1 expression or its activity, cause a reduction in the size of the vGPCR-induced tumors in mice^5^.

Regarding the role of HO-1 in the development of tumors, some results suggest that HO-1 can act as a cytoprotective enzyme, reducing the risk of developing some types of tumors. However, HO-1 is given a “dual” role. On one hand, it acts as a protective agent in healthy tissues but on the other hand it can act as an antiapoptotic and proangiógenic mediator.

In one way or another, and despite the fact that the role of HO-1 as a target of vGPCR is clear, we still do not know the mechanism that regulates the expression of its promoter. The transcription factors that bind to the HO-1 promoter are still poorly characterized, as the signal transduction pathways that regulate them.

The transcriptional regulation at the ARE site level is mainly controlled by Nrf2, a transcription factor of the leucine zipper type^6^. Under basal conditions, Nrf2 is found in the cytoplasm associated with a dimer of other proteins called Keap1^7^. This interaction stems regularly to Nrf2 towards the proteosome, so it becomes inactive and ends up being degraded. Faced with various inducers or stress agents, Nrf2 is released from Keap and is able to translocate to the nucleus where it acts as a transcription activator. In the nuclear interior, Nrf2 finds another associated protein, Maf, with which it forms a complex that binds to the ARE elements in the promoters of its target genes^7^.

In cells transformed by vGPCR it has been report the activation of at least three major signal transduction pathways MAPKS ERK2, JNK and p38^1,4^ Also the cascade of PI3K/Akt has been reported as activated by the viral oncogene. Activation of Akt would be dependent on PI3K and would have beta and gamma subunits of G proteins as intermediates^8^. Small GTP binding proteins of the Rho family, such as RhoA and Rac1, as well as alpha subunits of heterotrimeric G proteins increase their GTP loading in cells expressing vGPCR with their subsequent activation^4,5,9^. In particular we have shown that vGPCR induces HO-1 expression and cell transformation using a pathway that sequentially includes the Gα12 / 13 and RhoA proteins ^5,9^.

According to our previous results and in view of current global knowledge we hypothesize that Nrf2 acts as a key factor in the regulation of the promoter of HO-1 by vGPCR. We contribute new links that link vGPCR with the promoter of HO-1 highlighting the fundamental role of Nrf2 which not only in the regulation of HO-1 but also in tumorigenesis induced by vGPCR.

We have described that the loss of the ARE element (located at the −4 Kb position) results in a decrease in reporter activity indicating that this site is important in vGPCR mediated activation of HO-1 (Figure 1B), but since following deletions also show a decrease in reporter activation and that the shorter residual promoter retains some activation capacity, it is not ruled out that there are other important elements in the regulation of the HO-1 promoter. For example, we have seen that the deletion producing a 2.2 Kb fragment produces an increase in reporter activation indicating that between −3.8 Kb and −2.2 kb there is an element that negatively regulates this promoter and this observations awaits and deserves further analysis (data not shown).

We have determined the relevance of ARE sites on vGPCR-mediated activation. On one hand, we show that the deletion of a promoter region containing this sequence produces a decrease in the activation of the HO-1 promoter (Figure 1B) and on the other hand, that vGPCR is capable of activating a minimal promoter with ARE sites in tandem (Nrf2 ARE Luc) (Figure 2). Once this was demonstrated, we considered important to study the role of the transcription factor protein Nrf2. As shown in Figures 2 and 3, Nrf2 is key for the activation of the HO-1 promoter and the ARE sites.

Regarding the downstream elements of vGPCR it was described by our group and co-workers that vGPCR activates the HO-1 promoter through the small Gα12/13 G protein and RhoA ^5,9^. In this work, we demonstrated that both vGPCR and Gα12/13 activate ARE sites through RhoA and that they are not independent pathways since when co-transfected with a dominant negative of RhoA the increase induced by vGPCR and Gα12/13 was blocked (Figure 4B). This suggests that these elements are key in the activation of Nrf2.

vGPCR activates different transcription factors that control different genes. In our model, besides vGPCR has the ability to induce NF-κB through Rac1, and the HO-1 promoter has binding sites to this transcription factor, we show that the key transcription factor in the regulation of HO-1 mediated by vGPCR is Nrf2 and that its activation seems to proceed via RhoA and not Rac1, although a more thorough study is being currently followed in order to determine the nature of Rac involvement. All of this suggests a fine regulation by a complex network of proteins that are differentially activated in different models and that regulate different genes that orchestrate the biological response (proliferation, angiogenesis).

In this report, we have demonstrated that Nrf2 is able to recruit the transcriptional machinery and induce expression of a reporter gene in response, not only to vGPCR, but also in response to downstream elements (Figure 4A).

Another key aspect is the subcellular localization of Nrf2. In order to act as a transcription factor it is necessary that it can be translocated to the nucleus. We demonstrate here that both vGPCR and downstream elements are able to induce nuclear translocation of Nrf2 (Figure 6). This suggests that vGPCR is not only capable of inducing translocation to the nucleus of Nrf2 but that once bound to the ARE elements, Nrf2 is able to recruit the transcriptional machinery and initiate transcription.

There are numerous studies suggesting that phosphorylation of Nrf2 may contribute to its regulation. Nrf2 contains serines, threonines and tyrosines that can provide phosphorylation sites for various kinases^10^. For example, it has been shown that PKC can phosphorylate Nrf2 in serine 40 (Neh2) and disrupt the association between Nrf2 / Keap1 and thus promoting the translocation of Nrf2 to the nucleus ^11^. Keum et al have demonstrated that p38 can phosphorylate Nrf2, promote its association with Keap1 and thus prevent its nuclear translocation^12^.

Works from different groups have shown that vGPCR can activate different signaling pathways in different models. For example, Bais et al have demonstrated that in HEK293 cells vGPCR activates the JNK and p38 pathways^13^. In endothelial cells, vGPCR activates multiple pathways, including AMPc^14^ and AKT^8^. The activation of this pathway has also been described in NIH 3T3 cells. In our work, even using the same cell line we have not seen AKT activation (Figure 8A). We have shown that in our system, vGPCR activates the ERK1 / 2 and p38 pathways as shown in Figure 8 C-D.

We have shown that in our system vGPCR stabilizes Nrf2 and increases the phosphorylation levels of Nrf2. Furthermore, vGPCR activates the ERK1 / 2 pathway and has an effect on both Nrf2 transcriptional activation (Figure 9) and nuclear translocation (Figure 10). However, we have observed no effect on Nrf2 phosphorylation when we treated NIH 3T3 vGPCR cells with the MEK inhibitor PD98059 (Figure 11). This could be due, for example, to the fact that Nrf2 is being phosphorylated in several residues and that inhibition of the ERK pathway affects only certain amino acids among which Serine 40 is not found to recognize the antibody used herein. Another possibility that justifies this result would be that vGPCR activates the phosphorylation of Nrf2 by an ERK-independent pathway and that ERK is regulating some protein related to the import of Nrf2 to the nucleus. In this way, the inhibition of the ERK pathway would be impeding the translocation to the nucleus of Nrf2 without affecting its level of phosphorylation. We cannot determine with the techniques performed whether the increase of Nrf2 is due to an increase in the half-life of Nrf2 or if it is due to an increase in the expression of the Nrf2 gene However, our results clearly show the involvement of ERK2 as an intermediary between the vGPCR-Ga12/13-RhoA axis and nuclear translocation of Nrf2 as well as its transcriptional activation.

Studies published by Gjyshi et al have shown that de novo infection by KSHV leads to an increase in Nrf2 expression, an increase in the nuclear fraction of Nrf2 and an increase in phosphorylation levels of Nrf2^15,16^. In addition, they have demonstrated that the increase in Nrf2 stability is not due directly to the dissociation of Nrf2 from Keap1 but also increases the expression of Nrf2 and that increase in Nrf2 leads to an increase in HO-1. It is noteworthy that in those works they have use the complete genome of KSHV so that different proteins may be involved in the regulation of HO-1 but already with the complete virus is assigned a fundamental role to Nrf2. We, based on those works, can contribute that the effect observed by Gjyshi et al is probably due to vGPCR and this receptor might be responsible for the observed effects with de novo KSHV infection.

It is also important to note that when infected with the complete genome of KSHV many pathways begin to interact. For example, with regard to the increase of HO-1 we mention that KSHV increases the stability and phosphorylation of Nrf2, but it is also known that the BACH1 (repressor of HO-1 expression) mRNA is negatively regulated by viral miRNA MiR-K12-11^17,18^. This means that the expression of vGPCR and miR-K12-11 are two independent mechanisms that converge on the increase of HO-1.

By performing experiments in nude mice, we have shown that the tumorigenic effect of vGPCR is affected by the silencing of Nrf2. Cells that clearly show a decrease in Nrf2 expression by Western Blot were able to form tumors but with a markedly significant delay. It is interesting to note that immunohistochemical analysis of control and experimental groups have shown Nrf2 levels, but experimental groups have shown only residual levels. This data tells us about a positive selection of cells expressing Nrf2 produced in vivo on the population of injected cells and provides an extra data regarding the importance of expressing Nrf2 so that these cells can develop a tumor.

We knew that vGPCR was one of the key genes for tumor development induced by infection with KSHV. Our laboratory had previously reported that vGPCR was targeting the HO-1 promoter through the Gα12/13-RhoA proteins and that the development of these tumors was mediated by HO-1. We have also demonstrated that pharmacological inhibition or decreased HO-1 expression produced a decrease in tumor size^5^.

Throughout this work we have been able to deepen the study on the effects that vGPCR has on the expression of HO-1. We have shown that vGPCR not only activates Gα12/13 - RhoA but these proteins are signaling towards ARE sites present in the HO-1 promoter and that the transcription factor Nrf2 is key in this regulation. We have also show that vGPCR - Gα12/13 - RhoA is joined by MAPK ERK1/2 and that vGPCR also activates p38 MAPK. Regarding the role of ERK we have demonstrated that it has effects not only on the translocation to the nucleus of Nrf2 but also on the transcriptional activation of its transactivation domain. Finally, experiments in nude mice show that the tumorigenic effect of vGPCR is affected by the silencing of Nrf2 since mice injected with NIH 3T3 vGPCR - shNrf2 1 cells showed a delay in tumor development. All together our results point Nrf2 and its associated factors as a putative pharmacological target for controlling cell growth in cells transformed by KSHV oncogenes. We therefore aim our current efforts considering the Nrf2 system as a therapeutic target.

